# Exploiting epistasis to perturb the evolution of antibiotic resistance

**DOI:** 10.1101/738252

**Authors:** Marta Lukačišinová, Booshini Fernando, Tobias Bollenbach

## Abstract

New ways of curbing the ability of bacteria to evolve spontaneous resistance could mitigate the looming antibiotic resistance crisis. Progress toward this goal requires a comprehensive understanding of the key factors that contribute to resistance evolvability. Here, we present a systematic approach to identify cellular functions that affect the evolvability of resistance. Using a robotic lab-evolution platform that keeps population size and selection pressure under tight control for hundreds of *Escherichia coli* populations evolving in parallel, we quantified the effects of a genome-wide selection of pre-existing gene deletions on resistance evolution. Initial resistance of strains with gene deletions differed by more than tenfold but converged toward a hard upper bound for resistance during the evolution experiment, reflecting a global pattern of diminishing returns epistasis. We identified specific cellular functions that drastically curtail the evolvability of resistance; beyond DNA repair, these include membrane transport, LPS biosynthesis, and chaperones. Perturbations of efflux pumps prevented resistance evolution completely or forced evolution on inferior mutational paths, not explored in the wild type. We show that strong negative epistasis generally underlies these phenomena. The identified functions provide new targets for adjuvants tailored to block evolutionary paths to resistance when combined with antibiotics.

## Introduction

Bacterial resistance to antibiotics has become a major public health concern and a vibrant field of research (*1–3*). Strategies for countering the spread of resistance include the discovery of entirely new antibiotics and drug combinations (*4–6*). Recent work has increasingly focused on novel treatment schemes that minimize selection for resistance using drug cycling or combinations that exploit physiological or evolutionary interactions between drugs (*7–12*). Truly sustainable drug treatments require novel strategies that anticipate the evolutionary potential of pathogens and funnel them toward less evolvable genotypes or evolutionary dead ends. To this end, it is promising to identify genetic factors and cellular mechanisms that do not immediately increase a pathogen’s resistance but rather determine its ability to evolve (*13–16*).

The ability of different genotypes to spontaneously evolve drug resistance (here: “resistance evolvability”) can be determined by exposing them to equivalent selection pressures in evolution experiments and comparing the evolutionary outcomes (*17*). For a meaningful quantitative comparison, it is essential to control both the number of generations and the population size of the evolving genotypes (*18, 19*) – a characteristic which is achievable due to recent technological advances (*20, 21*). The outcome of such evolution experiments then depends on the key evolutionary determinants of the starting genotype, including its mutation rate and the distribution of fitness effects of resistance mutations.

The idea of interfering with resistance evolvability by altering the rate of acquisition of resistance by mutation or horizontal gene transfer is promising and has been investigated in depth (*13, 22–26*). Efforts to alter evolvability by other means, while equally promising, have received less attention (*16, 27*). Genetic differences commonly affect the fitness effects of new mutations (*28–31*). Such epistatic interactions can alter the effects of resistance mutations in different genetic backgrounds and potentially block mutational paths to drug resistance. For example, the gene coding for the transcriptional regulator AmpR opens a key path to resistance in *Pseudomonas*, since only strains that carry the gene rapidly evolve ceftazidime resistance by overexpressing beta-lactamase (*16*). Similar perturbations to global regulators could alter resistance evolvability more generally, since they completely change the expression state of the cell, and thus enable selection on newly expressed genes. In fact, perturbing any cellular function that interacts with a resistance mechanism – be it by interfering with its regulation, protein folding, localization, function, or degradation – could affect the cell’s ability to evolve. To discover mechanisms that determine resistance evolvability, a systematic investigation of resistance evolution starting from a diverse set of defined genotypes is needed.

Here, we report the discovery of targeted genetic perturbations that strongly affect antibiotic resistance evolution. We developed a robotic platform for high-throughput lab evolution, which tightly controls both population size and the selection pressure for drug resistance. Quantifying the evolvability of ~100 *E. coli* gene-deletion strains revealed a global trend of diminishing-returns epistasis: Genotypes that were initially more sensitive evolve resistance faster and converge to the same limit of resistance as initially resistant ones. We identified several specific genes and cellular functions that drastically affect – and, in some cases, entirely prevent – resistance evolution. These effects are largely due to epistatic interactions with common resistance mutations.

## Results

### An automated high-throughput evolution platform enables precise and reproducible measurements of resistance evolvability

To quantify the dynamics of resistance evolution for many different genotypes and replicates, we developed an automated platform that monitors the growth of hundreds of bacterial cultures while tightly controlling conditions and key evolution parameters (Fig. 1, Methods). Similar to the “morbidostat” setup (*21, 32*), the antibiotic concentration of each culture is periodically adjusted to maintain high selection pressure for antibiotic resistance over weeks. Every 3 to 5 hours, a dedicated robotic system dilutes and transfers cultures to new 96-well plates (Fig. 1A). In this transfer step, the volumes of medium, drug, and culture are individually tuned to keep each culture in exponential phase at 50% growth inhibition with defined population size right after the transfer (Fig. 1B-D, Methods). Keeping the cultures in exponential phase with vigorous shaking while continually adjusting the antibiotic concentration maintains a strong selection pressure, specifically for faster growth in the presence of the antibiotic and thus higher resistance (as measured by the IC_50_). In this way, fair comparisons of resistance evolution between hundreds of bacterial populations of the same size that undergo the same number of generations and experience the same clearly defined selection pressure become possible.

**Fig. 1.**
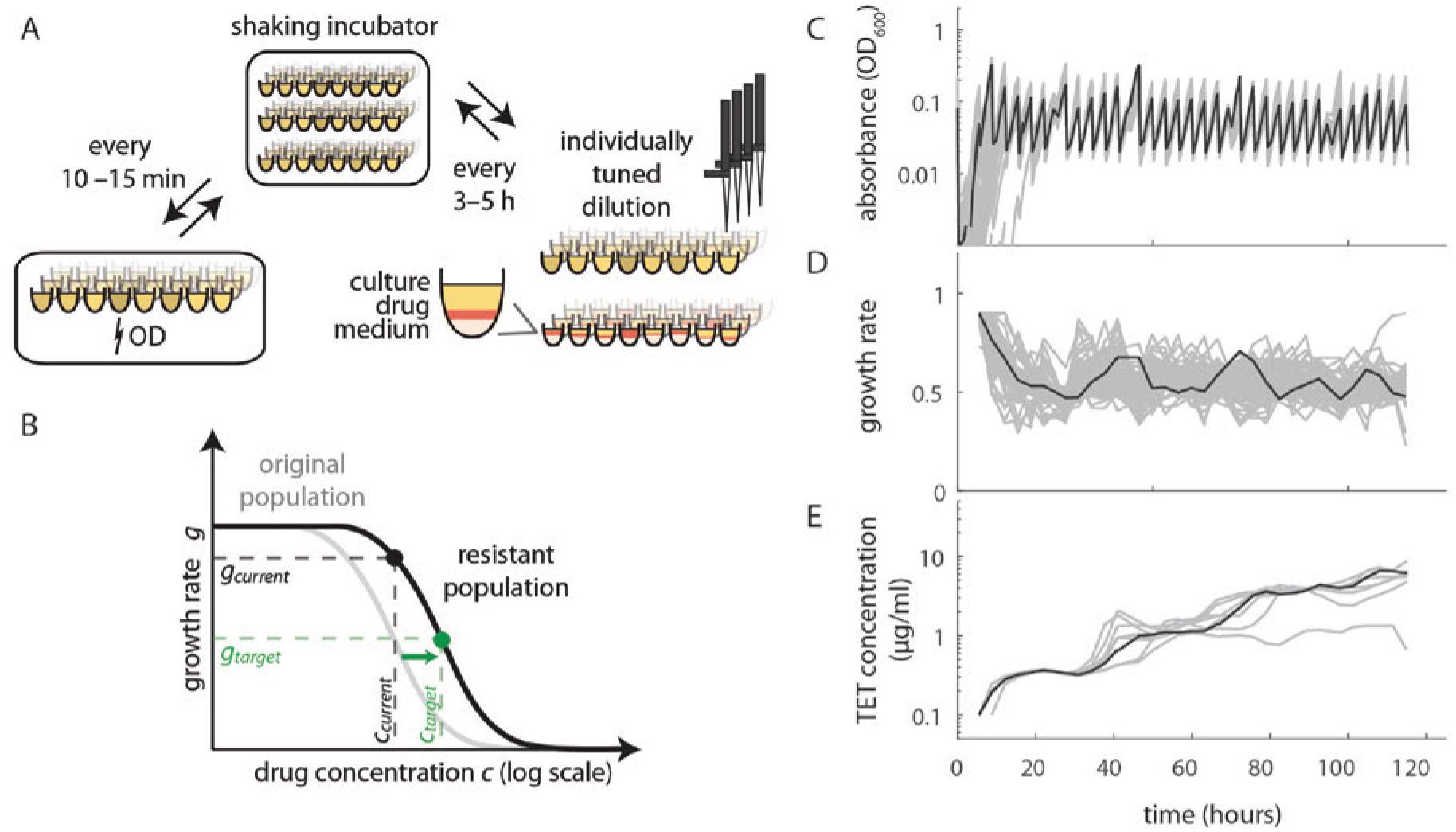
High-throughput lab evolution with controlled population size and selection pressure leads to repeatable fast evolution of antibiotic resistance. A) Schematic of lab evolution protocol. 96-well plates are shaken in an incubator, absorbance (OD) is measured every 10 to 15 min in a plate reader, and the cultures are diluted and transferred to new plates every 3 to5 h. At dilution, the volumes of culture, drug and medium are individually tuned based on the previous OD measurements, such that the OD after dilution and the growth rate is always close to the predetermined level (Methods). B) Schematic of dose-response curve in ancestral (gray line) and a resistant population (black line). At each dilution, the target antibiotic concentration *c*_target_ is calculated assuming that the the effect of resistance is merely is equivalent to reducing the concentration by a factor, i.e. a mere horizontal shift of the semi-logarithmic dose-response curve (*33*). The growth rate since the last dilution and antibiotic concentration in the given well (*g*_current_ and *c*_current_) then define the curve from which the new target concentration *c*_target_ is calculated. C) Background-subtracted OD values on log scale over the course of the experiment for all culture containing wells from a 96-well plate. The cultures are continuously in exponential phase. D) Growth rates fitted to the OD during the experiment. Values are normalized to the growth rate of the reference strain in no drug. In this case the reference strain is the parent strain of the Keio collection. E) Tetracycline concentration in the wells of the reference strain replicates during the experiment. The same replicate of the parent strain of the Keio collection is highlighted in black on all three plots (C-E). All values are from plate 1 of experiment M1 (Methods).

This lab evolution platform yields a precise and reproducible real-time measure of the resistance increase for each culture. Typically, resistance is measured by determining the minimum concentration needed to stop growth (MIC) or the concentration needed to inhibit growth by a certain factor, e.g. 50% (IC_50_). In our automated platform a feedback loop continually adjusts the antibiotic concentration to maintain 50% growth inhibition, therefore the antibiotic concentration in each well is a direct estimate of the IC_50_ (Methods). Indeed, we validated that this “on-the-fly” measurement of the IC_50_ agrees well with a standard IC_50_ measurement in a drug concentration gradient after the evolution experiment (Fig. S1). The strong selection pressure leads to rapid evolution: resistance in the wild type strain increases by 20-30 fold within a week (Fig. 2, Fig. S4) (*21*). Despite the fundamental stochasticity of evolution, the observed resistance increase over time is usually reproducible for replicates starting from the same genotype (Fig. 1E), in line with previous reports (*21, 33*). Whole-genome sequencing of evolved *E. coli* strains confirmed that the genes that mutate during the experiment are also largely reproducible (Methods, Fig. S2). For example, in the presence of the antibiotic tetracycline, the genes *marR, lon*, and *acrR* are often mutated at the end of the experiment (Fig. S2, Data S1), as previously observed (*34–36*). Typical evolved populations had three to four fixed mutations after ten days (Fig. S3). Taken together, our automated platform enables phenotypically and genotypically repeatable resistance evolution in high-throughput and is suitable for detecting perturbations that can alter resistance evolvability.

**Fig. 2.**
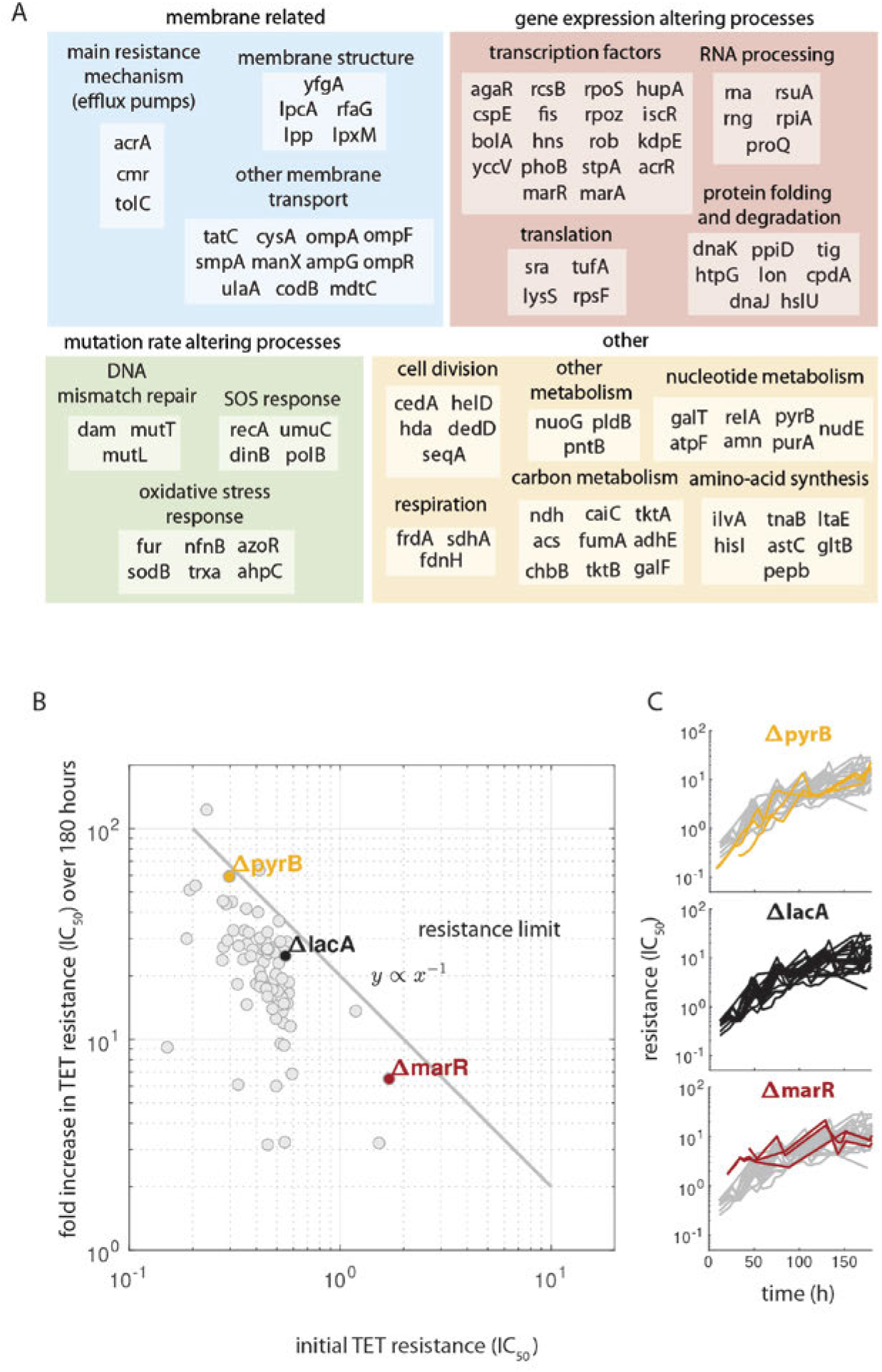
Resistance increases in diverse gene deletion strains reveal pattern of diminishing returns epistasis at the level of antibiotic resistance. A) Gene deletions chosen as ancestors for evolution experiments grouped by possible mechanism of evolvability alteration (large colored boxes) and specific cellular function (small boxes). B) Mean fold increase in resistance after 180 hours of evolution versus initial resistance for each deletion strain. The final and initial resistance measures for each replicate are the mean tetracycline concentrations over appropriate time intervals (12-24h for initial and 170-180h for final resistance). The line *y* ∝ *x*^−1^ indicates where the points would lie if all experiments reached the same final resistance irrespective of initial resistance. C) Resistance change over time replicates of three strains highlighted in B). Temporal evolutions of the reference strain (*ΔlacA*) are shown in gray for comparison. This figure uses data from experiment M2 in B) and combines data from M2 and M3 in C) (Methods).

### Antibiotic resistance evolution exhibits hallmark of diminishing returns epistasis

To investigate how diverse cellular perturbations affect resistance evolvability, we selected strains with genetic perturbations in a broad range of cellular functions from a genome-wide gene-deletion library (*37*) (Fig. 2A). The selection included both genes that we hypothesized to affect resistance evolvability and diverse other genes that were included to identify any general trends (Methods). For 13 out of 98 selected genes, we hypothesized that the impaired function has an effect on evolvability through mutation rate; these functions include DNA-mismatch repair (*38, 39*), SOS response (*13*), and oxidative-stress response (*23*). Further, we included deletions of known resistance genes such as drug efflux pumps (*40*), and porins (*41, 42*), together with several other membrane-related functions (Fig. 2A). Perturbations of functions that could systematically interfere with the expression and function of (yet unrecognized) resistance mechanisms were also included, in particular protein folding (*43, 44*), transcription factors, and membrane composition (Fig. 2A). The remaining 34 gene deletions were chosen to represent diverse cellular pathways that have negligible fitness costs and are expressed in rich medium (*45, 46*). The former is crucial to avoid selecting primarily for suppressor mutations of the gene deletion rather than for drug resistance (Methods). This broad selection of genetically different strains enables the discovery of general trends of resistance evolution and of cellular functions that lead to deviations from these trends.

We evolved the selected strains for tetracycline resistance and first identified general patterns that can explain the extent to which gene deletion strains evolved. A common pattern in evolution experiments is that identical beneficial mutations have weaker effects on fitter than on less fit backgrounds (*29, 30*). We observed a hallmark of such diminishing-returns epistasis at the level of drug resistance: Strains that were initially more sensitive underwent greater resistance increases during the experiment and effectively caught up with initially more resistant strains (Fig. 2B). Gene deletions often alter antibiotic resistance (*33, 46*); for tetracycline, the initial resistance levels of deletion strains varied by an order of magnitude (Fig. 2B). However, after 180 hours of evolution (about one hundred generations), these differences had largely evened out: Strains with an x-fold lower initial resistance (IC_50_) than the wild type tended to increase their resistance by x-fold more in the evolution experiment (Fig. 2B). This global pattern supports that there is a hard upper bound for spontaneous tetracycline resistance (“resistance limit”, Fig. 2B), which diminishes the possible resistance increases when approached.

### Perturbing efflux pumps drastically decreases resistance evolvability

Can cells be forced to deviate from this general trend of seemingly inevitable resistance? Noting that many spontaneous tetracycline resistance mutations directly relate to the overexpression of efflux pumps (*47, 48*), we reasoned that perturbing the composition or regulation of these pumps may affect resistance evolvability. Compromising efflux pumps sensitizes bacteria to various drugs (*46, 49*). According to the diminishing returns pattern (Fig. 2A), bacteria with perturbed efflux pumps are thus farther away from the resistance limit, and should evolve resistance faster. Alternatively, disrupting efflux pumps could effectively block this mutational path to resistance and force evolution to seek a different – likely less accessible – path. To discriminate between these two scenarios, we tested how perturbations of efflux pumps affect resistance evolvability.

Indeed, deleting the genes *acrA* or *tolC*, which code for components of the AcrA/B-TolC efflux pump, can essentially block resistance evolution. The most drastic effect occurred for *ΔtolC* in tetracycline, where we detected no increase in resistance in all seven replicates of the evolution experiment (Fig. 3B). This is striking because not only did this strain evolve under the same strong selection pressure for drug resistance at the same population size for the same number of generations as all other strains, but it was even five times more sensitive at the beginning of the evolution experiment (Fig. 3B) (*50*). We detected only one fixed mutation (a single base-pair substitution in the promoter of the *yhdJ* gene) in a single *ΔtolC* evolution replicate, corroborating the lack of adaptation; this idiosyncratic mutation appeared random and was unrelated to resistance (Data S1). Similarly, only one *ΔacrA* replicate out of five evolved tetracycline resistance, namely by overexpressing the homologous, rarely used AcrE/F-TolC pump, thus circumventing *acrA* loss (*51, 52*). The rapidity at which evolution found this alternative mutational path highlights the difficulty of perturbing resistance evolvability. The striking lack of resistance evolution for *ΔtolC* (Fig. 3B) likely reflects that TolC serves as an outer membrane channel for at least eight different efflux pumps (*53*), which can be disabled simultaneously. In sum, these results show how disrupting a specific cellular function can simultaneously sensitize cells to a drug and drastically slow resistance evolution.

**Fig. 3.**
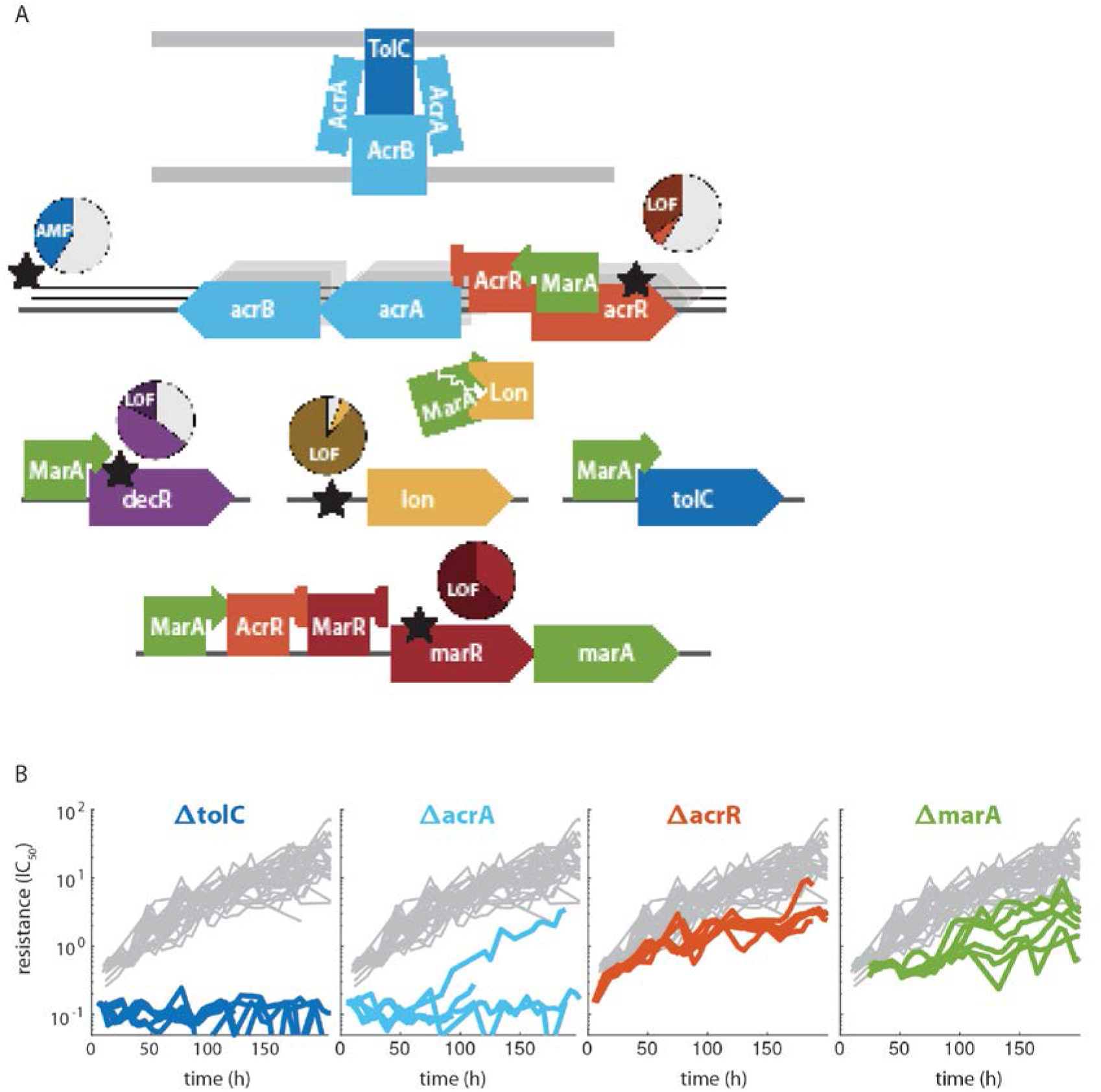
Perturbing multi-drug efflux pumps can drastically reduce the evolvability of tetracycline resistance. A) Schematic of AcrAB-TolC multidrug efflux pump in the membrane and genes that regulate it. Big arrows are genes, squares with pointing or blunt arrows are transcriptional activators or repressors respectively, and the Pacman shape is a protease. The five most common mutation loci are marked by stars; amplification of a genome region is shown as stacked black lines representing copies of DNA. Each mutation locus has a pie chart near it where colored slices represent the proportion of evolved populations started from the reference strain (*ΔlacA*) which gained a mutation in that locus during the experiment. Slices of a darker shade of the gene color represent the proportion of samples with predicted loss of function mutations in that locus. B) Resistance (IC_50_) over time for evolution experiments started from the *ΔtolC, ΔacrA, ΔacrR, ΔmarA* strains. All these genes are directly involved in the most common path to resistance shown in A. Gray lines show resistance over time for replicate experiments of the reference strain (*ΔlacA*).

We hypothesized that interfering with the regulation of efflux pumps while preserving their structural integrity provides additional ways to manipulate resistance evolvability. Indeed, several genetic perturbations of efflux-pump regulation significantly altered the rate of resistance evolution. Specifically, deleting *marA*, coding for a key activator of efflux-pump expression (*34, 54*) increases the initial sensitivity to tetracycline and slows subsequent resistance evolution, while not completely abolishing it (Fig. 3B). In contrast, deleting *marR*, coding for a repressor of *marA* (*55*) and therefore an indirect repressor of efflux pump expression, increases initial resistance but has no lasting effect on resistance (Fig. 2B), as expected from the diminishing returns pattern. Deleting *acrR*, coding for a repressor of the *acrA/B* operon and the *mar* regulon (*56*), shows increased initial sensitivity to tetracycline, likely because its coding region contains a MarA biding site that activates the *acrA/B* operon (*57*). Despite this initial sensitivity, resistance of the *ΔacrR* strain increases only modestly during the evolution experiment, by about 15-fold compared to 25-fold in the *ΔlacA* reference strain. Thus, deleting *acrR* is another way to decrease *acrA/B* expression and thus limit the attainable resistance. Taken together, we identified multiple cellular targets that slow resistance evolution by affecting efflux pumps, highlighting that even a single main mutational path to drug resistance can be targeted in several ways.

### Perturbing chaperones, lipopolysaccharide biosynthesis, DNA repair, and other genes can slow or accelerate resistance evolution

Beyond efflux pumps, we exposed additional cellular functions that slow resistance evolution when perturbed. In particular, deleting the gene for the Hsp70 chaperone DnaK (*58*) considerably slowed resistance evolution. The *ΔdnaK* strain evolved only 3-fold resistance while the reference strain reached more than 20-fold resistance gains in the same amount of time (Fig. 4). Furthermore, the *ΔdnaJ* strain which harbors a deletion of DnaK’s co-chaperone also exhibits slowed resistance evolution, albeit less extreme than *ΔdnaK* (Fig. S5). This observation is consistent with the notion that chaperones play a key role in evolution by affecting the conversion of genetic to phenotypic variability (*59, 60*). Perturbing different steps of the LPS biosynthesis pathway (*ΔlpcA* and *ΔlpxM*) led to over 4 and 2-fold lower levels of final resistance respectively (Fig. 4 and Fig. S5). Deletion of *tatC*, a gene involved in protein transport across the membrane, significantly slows resistance evolution (Fig. 4). This effect is possibly due to a lower mutation rate as this gene is thought to play a role in stress-induced mutagenesis (*61*). The molecular mechanisms underlying the effects of these genes on resistance evolution are unclear, highlighting the difficulty of predicting such evolvability modifiers and the importance of our systematic approach to exposing them. Together, these results support that multiple independent cellular functions determine resistance evolvability; perturbing these functions often defers resistance – exactly what is coveted in the clinic.

**Fig. 4.**
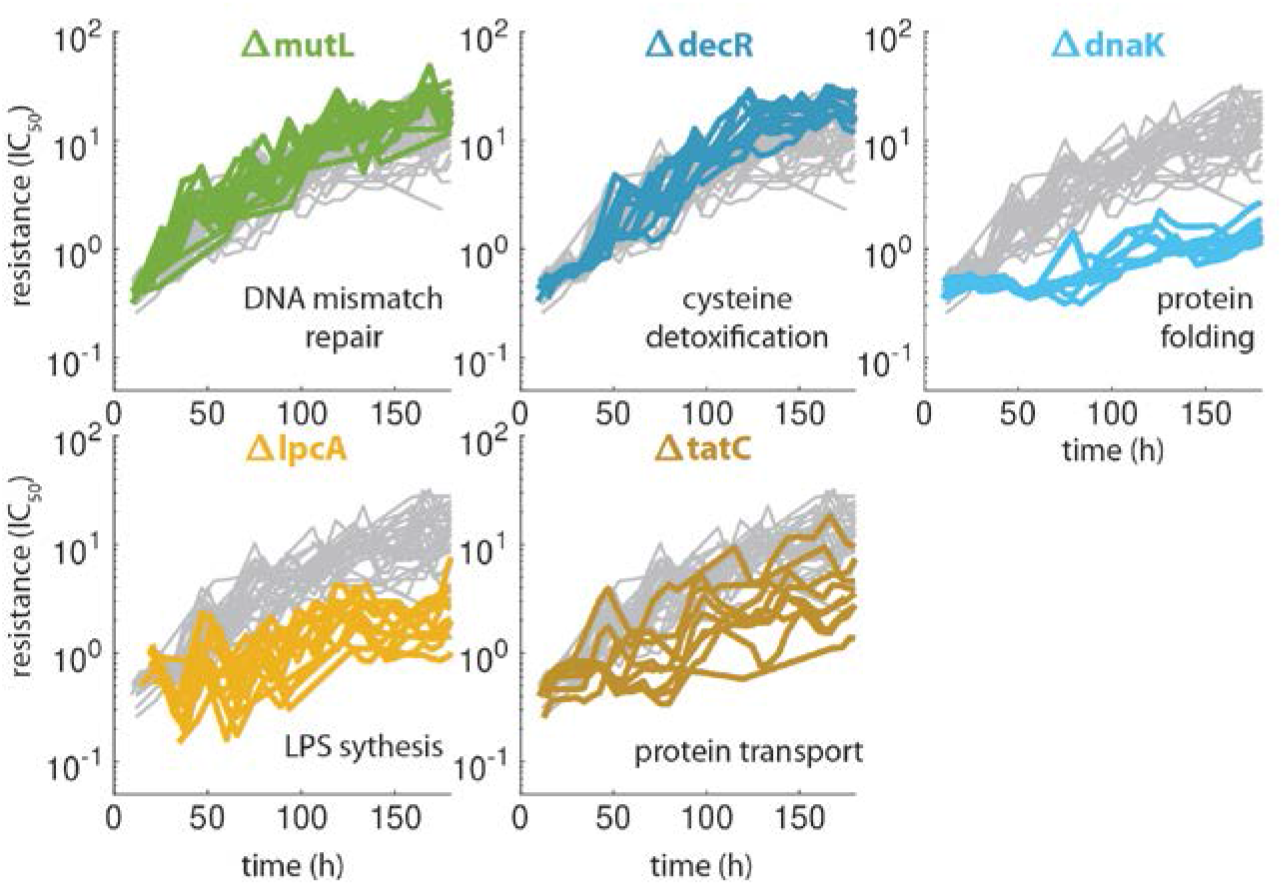
Diverse cellular functions, including chaperones, LPS biosynthesis, and DNA repair, affect resistance evolution. Resistance (IC_50_) over time as in Fig. 3 for several strains with deletions of genes not immediately related to efflux pumps which have a considerable effect on resistance evolvability. *ΔmutL* and *ΔdecR* show slightly increased evolvability, *ΔmutL* presumably due to the increased mutation rate. *ΔdnaK, ΔlpcA andΔtatC*, mutants in protein folding, LPS bioshynthesis and protein export respectively show decreased evolvability.

A few gene deletions also accelerate resistance evolution. Specifically, deleting *decR*, a regulator of cysteine detoxification (*62*), slightly accelerates resistance evolution even though it does not affect initial resistance (Fig. 4). We observed that a loss-of-function mutation in *decR* occurs reproducibly in the reference strain (Fig. 3A, Data S1). This is intriguing since the *decR* deletion alone does not increase resistance and suggests that loss of *decR* function amplifies the effects of spontaneous resistance mutations. A more straightforward way of accelerating resistance evolution are perturbations of DNA repair (*ΔmutL* in Fig. 4) that lead to mutator phenotypes with 100-fold increases in mutation rate (*38*). However, the effect is weak, suggesting that the occurrence of beneficial mutations is not rate-limiting for resistance evolution under our conditions. Overall, our observations support the notion that bacteria are more easily perturbed in ways that slow down resistance evolution rather than accelerate it. Importantly, this indicates a huge unexploited reservoir of candidate targets for choking resistance evolution.

To test if our results are specific to tetracycline or more generally applicable, we performed a similar evolution experiment with chloramphenicol. Like tetracycline, chloramphenicol targets the ribosome but the details of this interaction differ considerably (*63*). Whereas the evolution of tetracycline resistance seemed to level off within seven days, for chloramphenicol, we observed a steady increase even after ten days (Fig. S4, S6), confirming previous reports (*21*). Many mutations that fixed during the experiment overlapped with those observed for tetracycline, with additional mutations related to the MdfA efflux pump (Fig. S2) as previously described (*21, 33*). Despite these differences, the effects of specific gene deletions on evolution in the two drugs were remarkably similar (Pearson’s correlation coefficient r=0.77, p<10^−10^). In particular, the perturbations with the strongest effects were common to both antibiotics: Mutator strains (*ΔmutT* and *ΔmutL*) adapted faster while the *ΔtolC, ΔdnaK*, and *ΔmarR* strains adapted more slowly (Fig. S4). The accelerated evolution in mutator strains was clearer for chloramphenicol (Fig. S4). These results show that the effects of the cellular functions we identified are more general and do not just affect evolvability for one specific antibiotic in an idiosyncratic way, suggesting that hitting the same target can often modify resistance evolvability more broadly for different drugs.

### Changes in evolvability are largely caused by epistatic interactions with common resistance mechanisms

We hypothesized that many of the observed changes in evolvability are caused by epistasis between the gene deletions and common spontaneous resistance mutations. To test this hypothesis, we first combined our whole genome sequencing data for evolved strains with the resistance levels measured at the end of the evolution experiment. Based on these data, we built a simple linear regression model to estimate the benefit of each spontaneous resistance mutation (Methods). This model enabled us to identify deletion strains where these mutations fixed but had a different resistance benefit than expected (Fig. 5A). This analysis indicated magnitude epistasis, i.e. a quantitative change in the fitness effect of a mutation due to the presence of a different mutation (*31*), between the deletion and the acquired mutations: For example, in a *ΔdnaK* background, the same resistance mutations increased resistance by considerably less than in other strains (Fig. 5A). In extreme cases where the gene deletion completely nullifies the benefit of resistance mutations, these mutations would not fix in the evolution experiment (as observed for the *ΔtolC* strain, Fig. 3B). Thus, the strongest epistatic effects are not detectable by this approach.

**Fig. 5.**
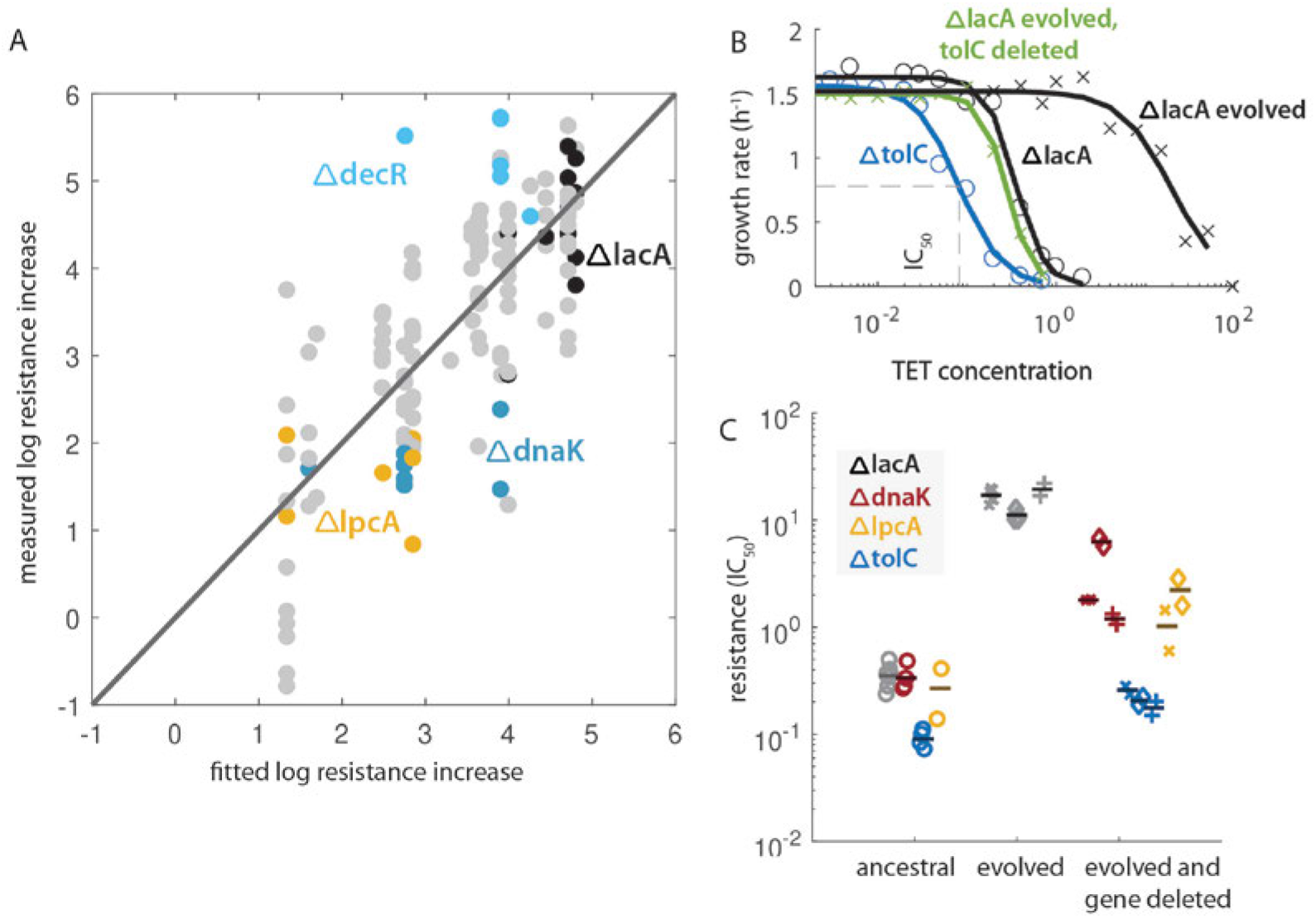
Epistasis between gene deletions and resistance mutations likely underlies altered evolutionary dynamics of deletion strains. A) Measured vs. fitted log resistance increase for each sequenced evolved population. For the prediction, the five most common mutations were considered (Methods). The black circles represent control (*ΔlacA*) strains. *ΔdecR, ΔlpcA* and *ΔdnaK* samples are highlighted to show that their effects systematically deviate from predictions. B) The dose-response curves for four strains. Black circles, blue circles, black crosses and dark blue crosses represent the ancestral *ΔlacA*, the ancestral *ΔtolC*, a clone isolated form an evolved *ΔlacA* population and the same clone with *tolC* deleted. A hill function is fitted to all four sets of measurements. The value for IC_50_ – the concentration at which the growth is half of the maximum – is shown with dashed gray lines. C) Resistance level (IC_50_) of the ancestral *ΔlacA* (reference, gray), *ΔdnaK* (red), *ΔtolC* (blue) and *ΔlpcA* (yellow) strains, three different resistant (evolved from *ΔlacA*) clones, and strains where *dnaK, tolC and lpcA* are deleted on the background of the respective resistant clones (Methods). Circles are used for ancestral strains, and crosses, diamonds and plus signs are used for three resistant clones isolated from three different evolutionary replicates starting from the reference *(ΔlacA)* strain. Horizontal lines represent the mean resistance value of each strain. The resistance decrease brought about by deleting *dnaK* on the resistant background is greater than on the sensitive background, showing epistasis between resistance mutations and *dnaK*. The same comparison is shown for the same resistant clones and *lpcA* and *tolC* deletions. Deleting *tolC* in the evolved strains brings their resistance to slightly below the initial resistance level of the reference strain.

To extend this analysis to extreme cases like *ΔtolC* and corroborate the central role of epistasis in resistance evolvability, we directly quantified epistatic interactions between specific gene deletions and common resistance mutations. We isolated clones from the *ΔlacA* reference populations evolved in tetracycline and deleted genes that modified the rate of resistance evolution in our experiments. We then measured the IC_50_ of the ancestral, evolved, and newly modified strains (Fig. 5B, Methods). Deleting *tolC* rendered the evolved reference strains even more sensitive to tetracycline than the reference strain was at the beginning of the experiment (Fig. 5C). The deletion of *dnaK* and the deletion of *lpcA* also sensitized the resistant strains, albeit only partially (Fig. 5C). These results expose epistatic interactions between resistance mutations and gene deletions identified in our large-scale search for evolvability modifiers, which are consistent with their slower resistance evolution. More quantitatively, the extent of epistasis mirrored the observed differences in resistance evolvability for *ΔtolC* and *ΔdnaK*, respectively (Fig. 3 and 4). Together, these results highlight the potential of exploiting epistatic interactions for evolvability modification as a promising strategy to restrain resistance evolution.

## Discussion

We presented a systematic analysis of the effects of targeted genetic perturbations on antibiotic resistance evolution. Using automated high-throughput evolution experiments, we identified general patterns guiding resistance evolution and specific cellular functions that affect resistance evolvability most drastically. We established a high-throughput experimental evolution platform that keeps hundreds of cultures in parallel in exponential phase under controlled selection pressure. This platform allows precise detection of adaptation rates over a wide dynamic range; it enables quantitative investigations of evolvability for diverse microbes, including the most worrisome pathogens, and other stressors than antibiotics.

Resistance-enabling genes like *tolC*, *dnaK*, *lpcA*, which drastically slow resistance evolution when deleted (Fig. 3 and 4), are candidate drug targets for a new strategy in which antibiotics are combined with compounds that do not lead to immediate synergy but slow down, or perhaps even prevent, resistance evolution in the long term. Antisense oligomers (phosphorodiamidate morpholino oligomers, PPMOs) are a promising way to inhibit the expression of a broad range of targets (*64*) – an approach that has been successfully used for genes encoding efflux pumps and leads to antibiotic hyper-sensitivity (*50*). Targeting efflux pumps would also be possible with efflux-pump-inhibiting molecules (65–67) or phages which require TolC for entry (*68*). TolC, which has received considerable attention (*50, 69, 70*) is a particularly interesting target in this context: A TolC-inhibitor would strongly synergize with antibiotics like tetracycline and chloramphenicol (*50*) while at the same time slowing resistance evolution. Thus, this strategy may solve the catch-22 that the coveted synergistic drug combinations tend to accelerate resistance evolution (*71*). Discovering inhibitors of the newly identified evolvability modifiers could reinvigorate old drugs and, at the same time, put their use on a sustainable future trajectory.

The detailed molecular mechanisms behind the altered evolutionary dynamics of *ΔdnaK, ΔlpcA, ΔtatC* and *ΔdecR* remain unclear. Chaperones such as DnaK were famously proposed to affect evolution by buffering the phenotypic effects of mutations (*43, 60, 72*). However, there is also evidence that chaperones may enhance the phenotypic effects of spontaneous mutations (*73*). In the case of efflux-pump related resistance, a possible link is that TolC is a predicted client of DnaK (*74*). In addition to sensitizing bacteria to antibiotics, perturbing membrane composition via LPS biosynthesis could also influence membrane permeability for antibiotics and thus interfere with the effects of resistance mutations. However, there is evidence that just perturbing the assembled LPS layer does not change efflux pump activity (*75*). The mechanisms underlying the faster adaptation of the *ΔdecR* strain (Fig. 4) and the frequency of spontaneous mutations in this locus remain to be elucidated. DecR was recently shown to be a repressor of only one operon, which is involved in L-cysteine detoxification (*62*); *decR* is upregulated by MarA and TolC is involved in L-cysteine transport (*76*). Therefore, the resistance mutations affecting these genes may also affect DecR and L-cysteine levels, introducing a cost or sensitivity, which can in turn be alleviated by specific mutations in this regulator. Elucidating the molecular mechanisms underlying evolvability modification is an exciting future challenge.

Even when resistance-enabling genes like *tolC* or *dnaK* are deleted, evolution might ultimately find ways to increase resistance, but this can take orders of magnitude longer. A powerful application would be to combine an antibiotic with inhibitors for several of the key resistance-enabling genes identified using the approach presented here. In this way, even less-common paths to resistance could be blocked. The probability of circumventing these blocks by mutation may become prohibitively low. Thus, in practice these combinations could be a key step toward finally casting Ehrlich’s elusive “magic bullets” after trying for over 100 years. This work lays the foundation for realizing this vision in the future by extending our approach to the genome-wide scale, to other drugs, and to other organisms, in particular to the most relevant pathogenic microbes.

## Acknowledgments

We thank Erdal Toprak, Nassos Typas, Marjon deVos, and Marcin Suskiewicz for critical comments on the manuscript and members of the Bollenbach group for fruitful discussions.

## Author contributions

ML and TB conceived the study and designed the experiments. ML performed the experiments and analyzed the data. BF constructed strains and performed the experiment shown in Fig. 5B,C. ML and TB wrote the manuscript.

## Supplementary Material

### Materials and Methods

#### Strains, media, reagents and antibiotics

Cultures were grown in LB medium from Sigma Aldrich (#L3022). For PCR reactions GoTaq G2 DNA Polymerase (Promega #M7845) or Q5 high fidelity Polymerase (New England Biolabs #M0491S) were used.

All strains originated from isolated clones (plated on solid LB, picked, regrown overnight in LB and frozen in 15% glycerol) from the Keio collection (Baba et al., 2006) with kanamycin cassette included in the locus of the deleted gene.

Tetracycline stock solutions of 7mg/ml or 10mg/ml were prepared by diluting tetracycline hydrochloride powder (Sigma Aldrich # T7660) in 83% ethanol at room temperature. Chloramphenicol stocks of 10mg/ml were prepared by diluting powder (Sigma Aldrich #C0378) in 99% ethanol. Kanamycin stock was made from kanamycin sulfate powder (Sigma Aldrich #K4000). All antibiotic stocks were stored at −20°C.

#### Whole-genome sequencing analysis

Whole genome sequencing was performed for 380 samples altogether as listed in Data S1. For all evolved population samples, the ancestral clone was also sequenced and its mutations analyzed Data S1), to distinguish clearly between mutations acquired before and during the experiment. Genomic DNA was purified directly from thawed glycerol stocks using the GenElute 96 Well Tissue Genomic DNA Purification Kit (Sigma-Aldrich # G1N9604). Library preparation, multiplexing, and sequencing were performed by LGC Genomics GmbH. The samples were sequenced on an Illumina NextSeq500 V2 (paired-end sequencing, 150bp read length, ~230-fold coverage on average, but ranging from ~70-to ~800-fold due to the multiplexing protocol). Sequencing data were analyzed using Breseq (*74*) (Version 0.32.0). Reads were aligned to the deposited Keio parent reference (Accession: CP009273) using Bowtie2. The mutations identified by Breseq were manually inspected for false positives; all validated mutations are listed in Data S1. Even though the samples were expected to be heterogeneous (they were not isolated clones), the “clonal” mode of Breseq was used. Therefore, the mutations detected only represent fixed mutations. Amplifications were noted if the coverage of a multi-genic region exceeded twice the average coverage of that sample. Since an IS insertion in the *lon* promoter region was often among the “unassigned new junction evidence” but at very high frequency, this type of mutation was assumed to be fixed if the frequency exceeded 90%. For each evolved sample, we validated that the intended gene deletion is present. If any reads in the deletion locus were present, which would suggest cross-contamination with another strain, the sample was excluded from the analysis, as indicated in Data S1.

#### Automated experimental evolution

The selected deletion strains from the Keio collection (*37*) as listed in Table S1 were all streaked for single colonies and clonal cultures frozen with 15% glycerol at −80°C. The glycerol stocks were used to assemble the starting 96-well plates for the evolution experiment. Each plate had at least 12 empty wells, which were filled with sterile growth medium, and handled like all other wells throughout the experiment to monitor cross-contamination. Replicates of the same ancestor, if on the same plate, were placed far from each other to avoid cross-contamination which would not be detected by genotyping. Every plate contained at least two replicates (and usually more) of the control strain (either the BW25113 parent strain of the Keio collection or the *ΔlacA* strain). The *ΔlacA* strain was used as reference in later experiments instead of the BW25113 to ensure that any difference between the strains is not due to the presence or absence of the kanamycin cassette.

The automated evolution protocol was carried out four times using a Tecan Freedom Evo 150 liquid-handling platform. The specific differences between the runs of the experiment are given in Table S2. The 200μl cultures were kept in LB rich medium in 96-well plates (Nunc, transparent flat-bottom) in a shaking incubator (Liconic Storex, 30°C, >95% humidity, 720rpm). Continual shaking and rich growth medium ensured that the cultures were homogeneous and not under oxygen or nutrient limitation. Every 10 to 15min, each plate was transferred to a plate reader (Tecan Infinite F500) using a robotic manipulator arm (RoMa) and the absorbance (OD at 600 nm) was measured. Every 3-5h, the cultures were transferred to new plates. They were not diluted in the same plates to avoid biofilm formation and due to large errors in volumes left in the wells after pipetting out most of the culture. The new plate was filled in three steps. First, pure LB medium (*v*_med_) was pipetted, then medium with antibiotic (*v*_ab_) and last the culture (*v*_culture_) from the previous plate. Each culture had its own dedicated 200 μl disposable tip, which was washed in ethanol after every dilution. LB medium and antibiotic stock were multi-pipetted into the new plates using 1000 μl tips. All tips were exchanged once a day. The reservoirs with media had lids that were taken off using the RoMa arm just before usage.

Every day of the experiment, the penultimate plates of that day were left in the incubator to grow out: the next day 70μl of 50% glycerol was added to each well and the plates frozen at −80°C. Fresh antibiotic stocks and medium reservoirs were provided. There were always two concentrations of antibiotic stocks available, approximately 10-fold apart, the protocol always only chose one of the available stocks to pipette from. The concentrations of the antibiotic stocks were chosen each day depending on how resistant the populations had become.

Every 3 to 5 hours the cultures from each plate were transferred to new plates using the Air LiHa robotic pipetting head. The appropriate volumes of culture, medium and antibiotic to use were calculated at each dilution step and for each culture using a custom Python script based on the OD values obtained since the last dilution. The growth rate was obtained from 18 consecutive OD measurements by obtaining the slope of the least-squares linear fit (`numpy.polyfit` function) to the log2 of those background subtracted OD values which were between 0.01-0.1. All growth rates were normalized to the growth rate of the reference strain in the absence of antibiotic (1.7 doublings per hour). The volumes were calculated separately for each well using the last OD measurement (*d*), normalized fitted growth rate (*g*_current_), concentration of antibiotic stock (*c*_stock_), current antibiotic concentration in the well (*c*_current_) and Hill coefficient of the dose response curve *n*_TET_ = 1.8, *n*_CHL_ = 2.4 (*33*) in order to reach the target OD (*d*_target_ = 0.01), growth rate (*g*_target_ = 0.5) and total volume (*v*_total_ = 200) according the equations:

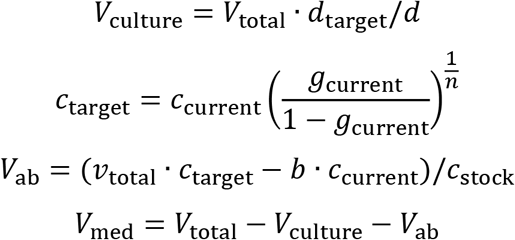

We took several precautions to deal with atypical input values. If the concentration *c*_current_ is zero, *c*_target_ is set to a default concentration of 0.1 μg/ml for tetracycline and 0.5 μg/ml for chloramphenicol, which are values lower than the IC_50_ of the most sensitive strains in our selection. If the sum of squared residuals from the fit to obtain the growth rate is greater than 0.8, which was empirically chosen to reflect a failed growth rate fit, then *c*_target_ is set to *c*_current_. If the measured normalized growth rate is larger than 0.9, it is set to 0.9 to avoid very large or undefined values for *c*_target_ due to the sigmoidal shape of the dose-response curve. If the calculated volume *V*_ab_ is smaller than 5μl, *V*_ab_ is set to zero (only medium is used to dilute the culture) and concentrations are updated accordingly. *V*_culture_ is capped at 140μl, to assure accurate aspiration from the small 200μl culture. There were two available reservoirs of antibiotic stocks, the higher concentration was only used if, for the lower stock concentration *V*_ab_ > *V*_total_ – *V*_culture_.

#### Resistance measure

The “on-the fly” resistance (IC_50_) measure for a particular culture is the antibiotic concentration in the well at that time. The concentration was updated at every dilution. For all plots of resistance over time (Fig. 2–4) and all “initial” and “final” resistance measures, only those time points where the growth rate after that particular dilution was close to half-inhibited (between 0.3 and 0.7 of the maximum growth rate of the reference strain) were considered.

#### Regression model of mutational effects

The regression model has two major assumptions. First, different mutations in the same locus provide the same resistance benefit and second, the effects of mutations on resistance are additive on a log scale, i.e. each mutation brings a fixed relative resistance increase irrespective of which other mutations are present. Assuming this is true, the log resistance y can be expressed as a linear model:

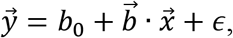

where 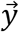 is the log of the increase in resistance observed for the individual evolving populations, *b*_0_ is a fitted coefficient corresponding to the resistance increase common to all evolved populations not predicted by the five most common mutations, 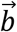 is the vector of fitted coefficients which correspond to the effects of the individual mutations, 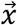 is a vector of 1s or 0s determining the presence or absence of that particular mutation in the given evolved population.

Mutations from all sequenced samples evolved in tetracycline which passed contamination and quality control were included in the analysis Data S1. The function `fitlm` (Matlab R2016b) was used. The predicting variables were the presence and absence of mutations in the five most commonly hit genes (*marR, lon*, efflux-pump amplification, *acrR, decR*), the fit parameters were the multiplicative effects of mutations in those loci, and the response variable was the log resistance increase over the course of the experiment. The predicted mutational effects of the five most common mutations are given in Table S3.

#### Generation of double deletion mutants

Before introducing the kanamycin cassette into the evolved *ΔlacA::kan^R^* strains to produce double-deletion mutants, clones were picked from LB agar plates with 25 μg/ml kanamycin-sulfate and 10 μg/ml tetracycline hydrochloride and their growth rates were compared with the evolved strain. The growth rate was determined in a dose-response assay as explained in “Dose-response measurements”. Clones with resistance level similar to the evolved population were subjected to P1-phage transduction (for *dnaK* and *tolC* deletion) or lambda-red recombineering (for *lpcA* deletion).

Prior to the P1-phage transduction, the FRT-flanked kanamycin cassette has been removed from the evolved *ΔlacA* strains with the plasmid pCP20 and selection on LB agar with 100 μg/ml ampicillin and with 25 μg/ml kanamycin. Afterwards the double-deletion mutants were created by transferring the respective alleles (*ΔdnaK, ΔtolC, ΔlpcA* and *ΔlacA*) from the Keio collection into *ΔlacA* evolved strain using the standard P1-phage transduction protocol (*75*). The genotype was verified after P1 transduction with PCR (Table S4).

To delete the *lpcA* gene in the evolved *ΔlacA* strains, the chromosomal gene *lpcA* was targeted with lambda-red mediated homologous recombination, due to inefficient P1 infection of the *ΔlpcA* strain. A PCR product containing the kanamycin cassette flanked by FLP recognition target sites and 50 base pairs homologies to adjacent chromosomal sequences, as described elsewhere (*37*)(Table S4), and 20 bp homology to the plasmid pKD13, were amplified using Q5-HF-polymerase (NEB). The PCR product was purified using a standardized PCR clean-up kit (Promega #A9282) and electroporated into evolved *E. coli* BW2511 *ΔlacA::kan^R^* with the recombineering plasmid pSIM19. The transformed cells were selected for kanamycin (25 μg/ml) and the presence of the PCR product was confirmed by colony PCR (Table S4)

#### Dose-response assay

Strains were grown overnight at 30°C in LB broth without any antibiotics for 20 hours. The growth rate of the double- and single-gene deletion mutants with and without tetracycline were determined at OD_600_ using the Biotek plate reader Synergy H1. The overnight culture was diluted 1:1000 in all assays. The cell growth was observed for 25 hours at 30°C.

The Hill function fits were obtained by fitting the function

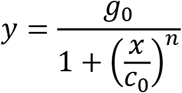

to the growth rate measurements using the function fit (Matlab R2016b). *g*_0_ is the fitted maximum growthrate (or the growth rate without drug), c0 is the fitted IC_50_ and *n* is the fitted dose sensitivity (*33*).

#### Growth rate fits

Unless specified otherwise, growth rates are determined as the slopes of a linear fit to the log2 background subtracted OD values. Only those OD values which lie between 0.015 and 0.1 (after background subtraction) and only the time window from when the values first cross 0.015 until they reach 0.2 were considered. The function fit (Matlab R2016b) was used.

**Fig. S1.**
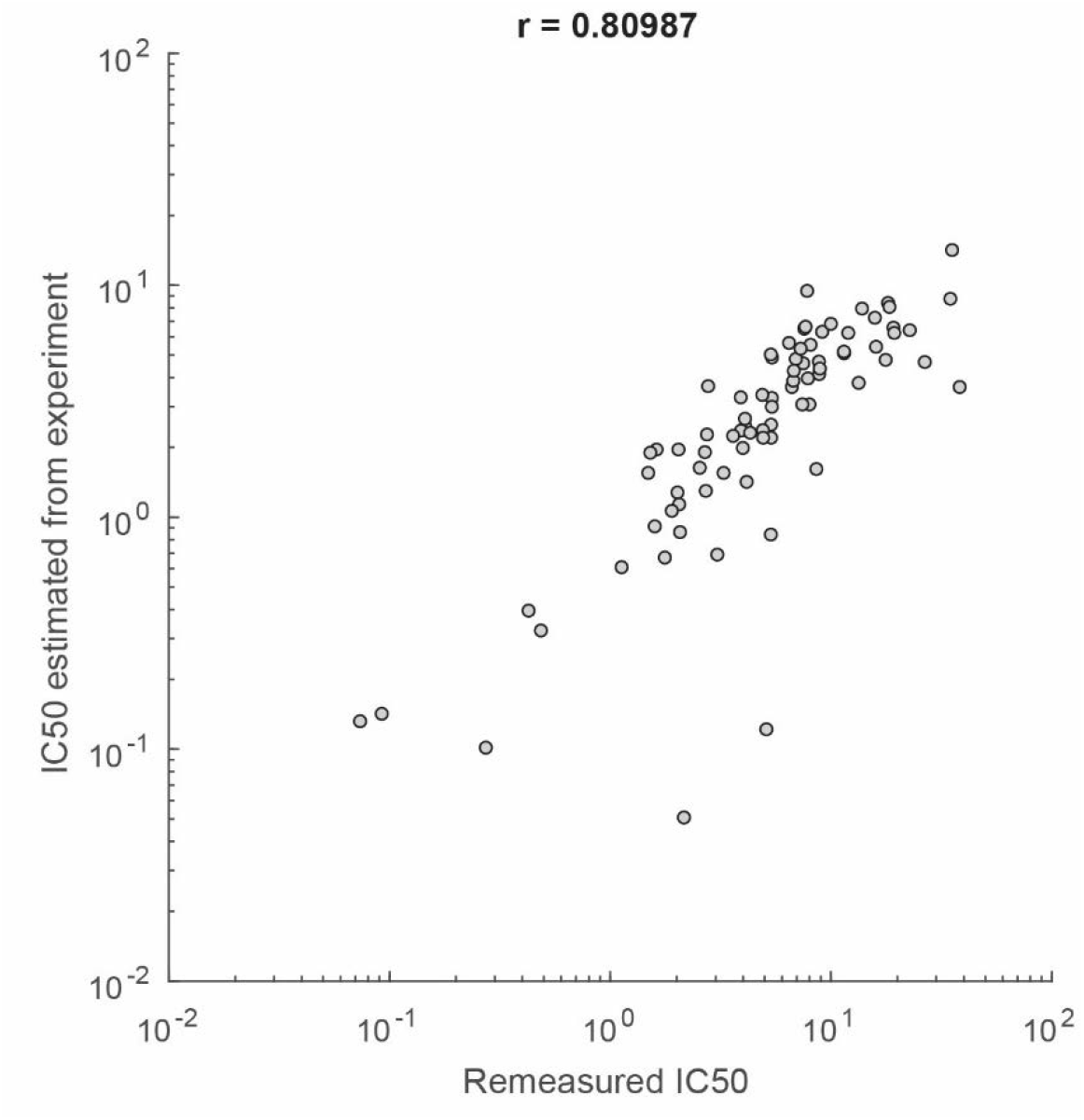
Antibiotic concentration in well during evolution experiment agrees with conventional IC50 measurement. The concentration of tetracycline in the well at the end of the experiment for many wells is plotted against the fitted IC50 values from measuring the growth rate of the same populations in a wide range of tetracycline concentrations (Methods). Pearson’s correlation coefficient calculated from the log values of the two measurements is given in the title of the plot.

**Fig. S2.**
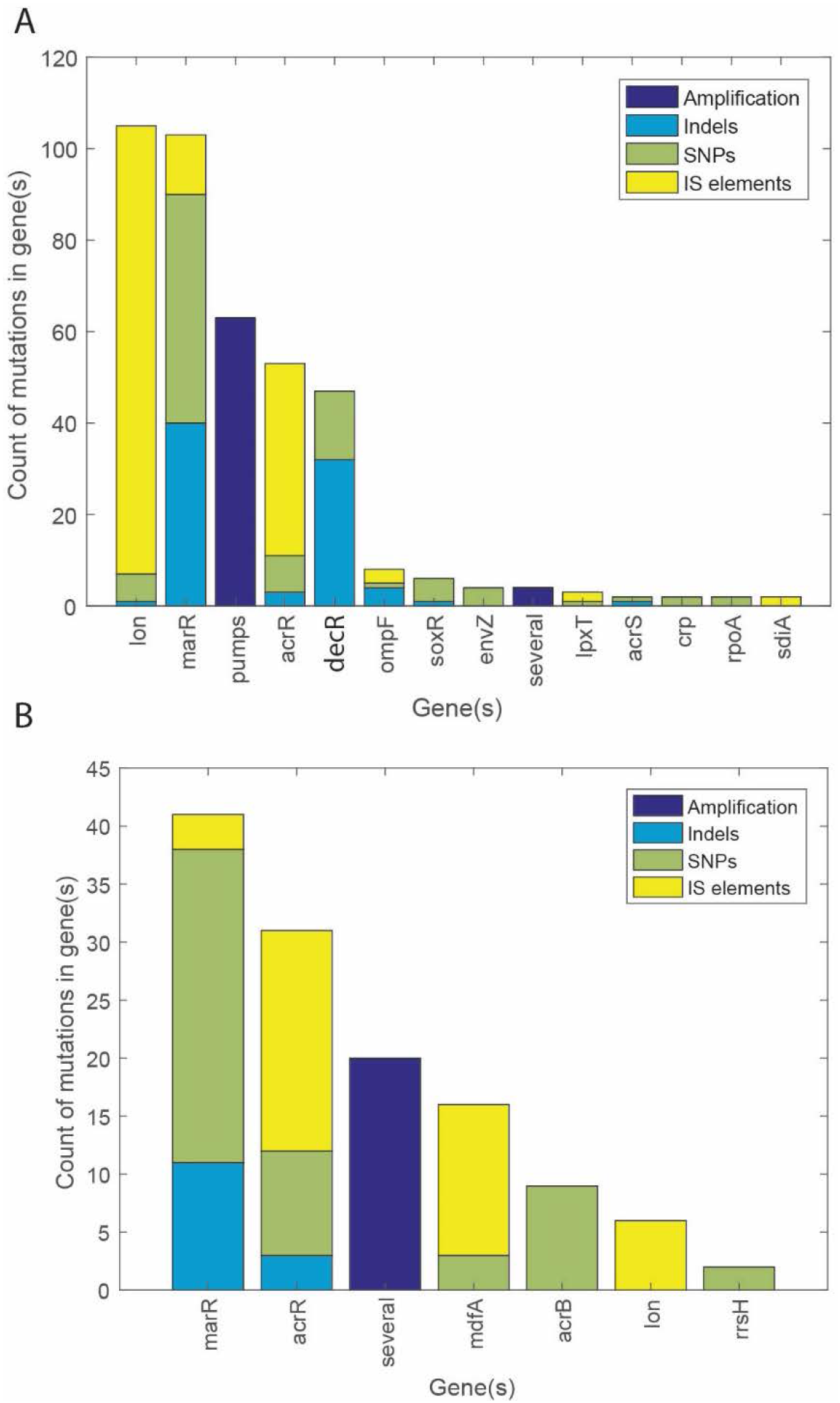
Mutations reproducibly occur in a small number of loci during evolution in tetracycline and chloramphenicol. A) Counts and types of fixed mutations found in populations evolved in tetracycline grouped by gene locus where they occurred. Only genes which were hit at least twice in our dataset are shown. The label “pumps” denotes an amplification of the region of the acrA/B operon identified from a more than two-fold increase in coverage compared to the mean (Methods). The label “several” denotes an amplification of a different region spanning several genes. The five most common mutations represent 84% of the total mutations identified. B) The same chart as in A) for mutations identified in populations evolved in chloramphenicol.

**Fig. S3.**
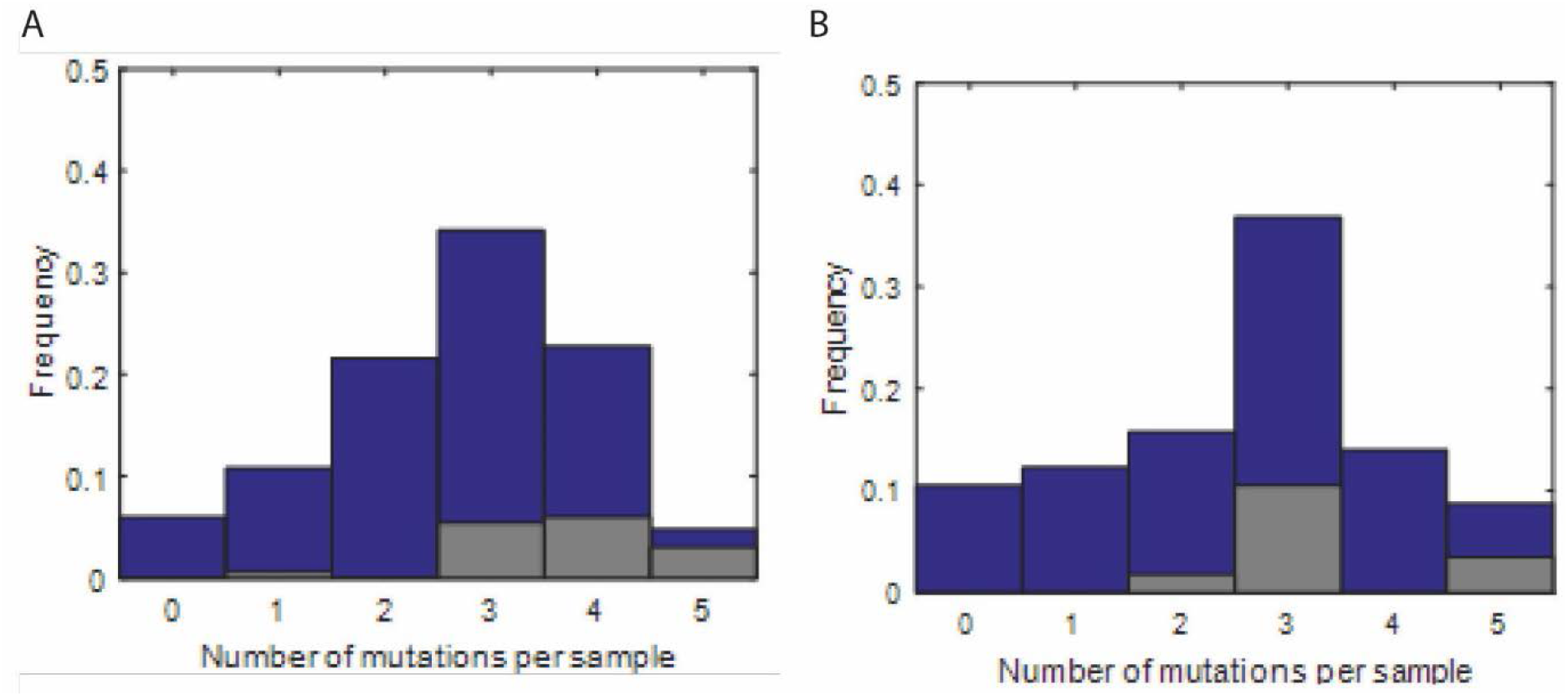
Number of identified fixed mutations in evolved populations. A) Histogram of the number of mutations detected in each evolved sample. The gray bars represent the number of mutations found in reference strains: the Keio parent strains or the ΔlacA strain. The typical number of fixed mutations for reference strains is 3 or 4. Since the selection sent for sequencing was biased toward slowly evolving strains, the overall counts are biased to fewer mutations. B) The same chart as in A) for the counts of mutations identified in populations evolved in chloramphenicol.

**Fig. S4.**
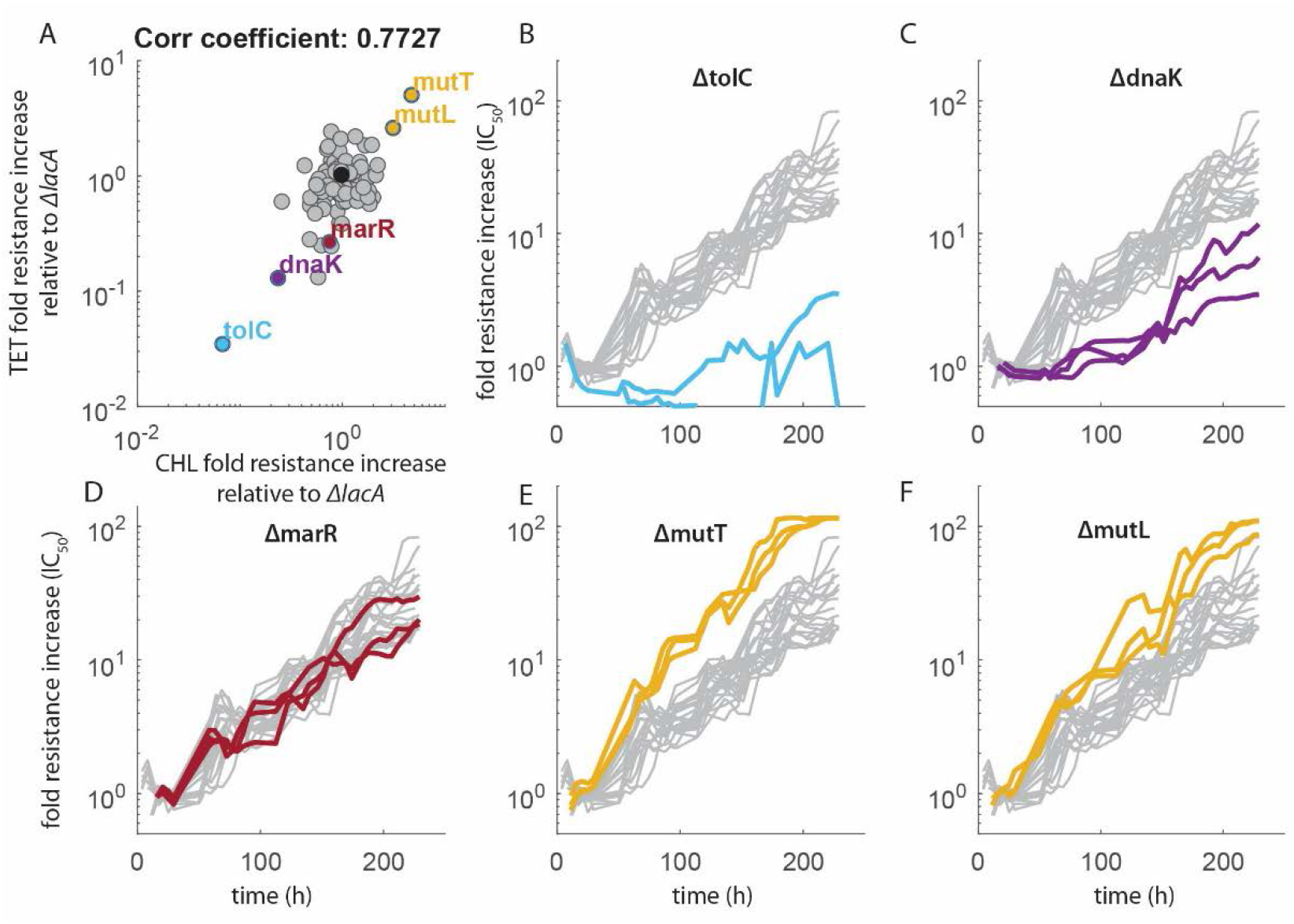
Strain specific changes in tetracycline and chloramphenicol evolvability are correlated. A) Fold increases in resistance normalized to the mean fold increase in resistance for the reference strain (*ΔlacA*) shown in black. The correlation coefficient is given in the title of the plot. B-E) Examples of resistance increase over time for chloramphenicol for strains highlighted in the upper left plot. The apparent plateauing of the resistance for *ΔmutT* is due to the fact that the concentration needed to keep the strains inhibited simply reached the maximum stock concentration used on that day.

**Fig. S5.**
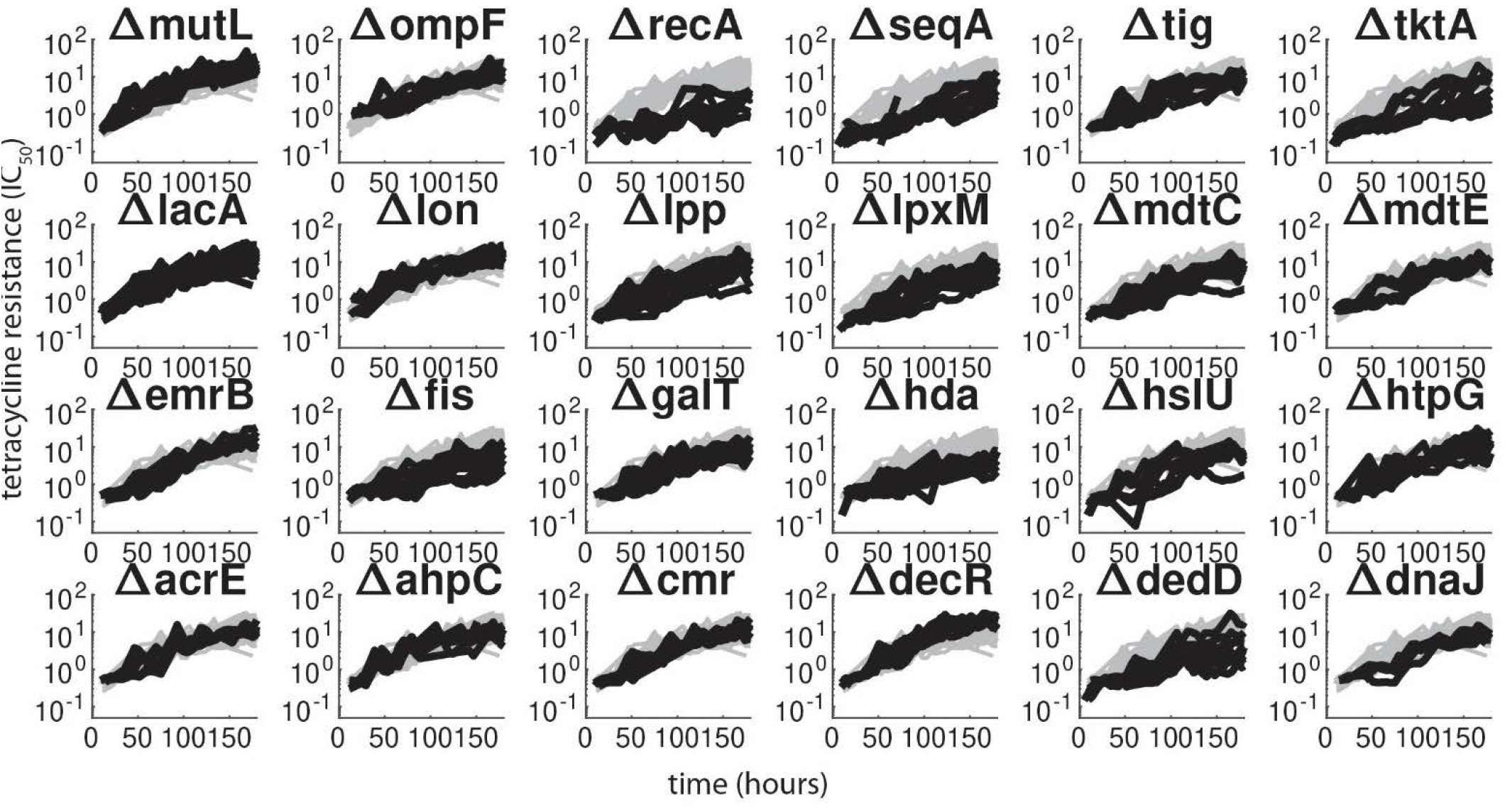
Resistance increases over time in tetracycline for additional ancestral strains. Resistance (as measured by the antibiotic concentration in the well) over time for all deletion strains for which more than 3 evolutionary replicates in tetracycline were done and which were not shown in Figures 2–4.

**Fig. S6.**
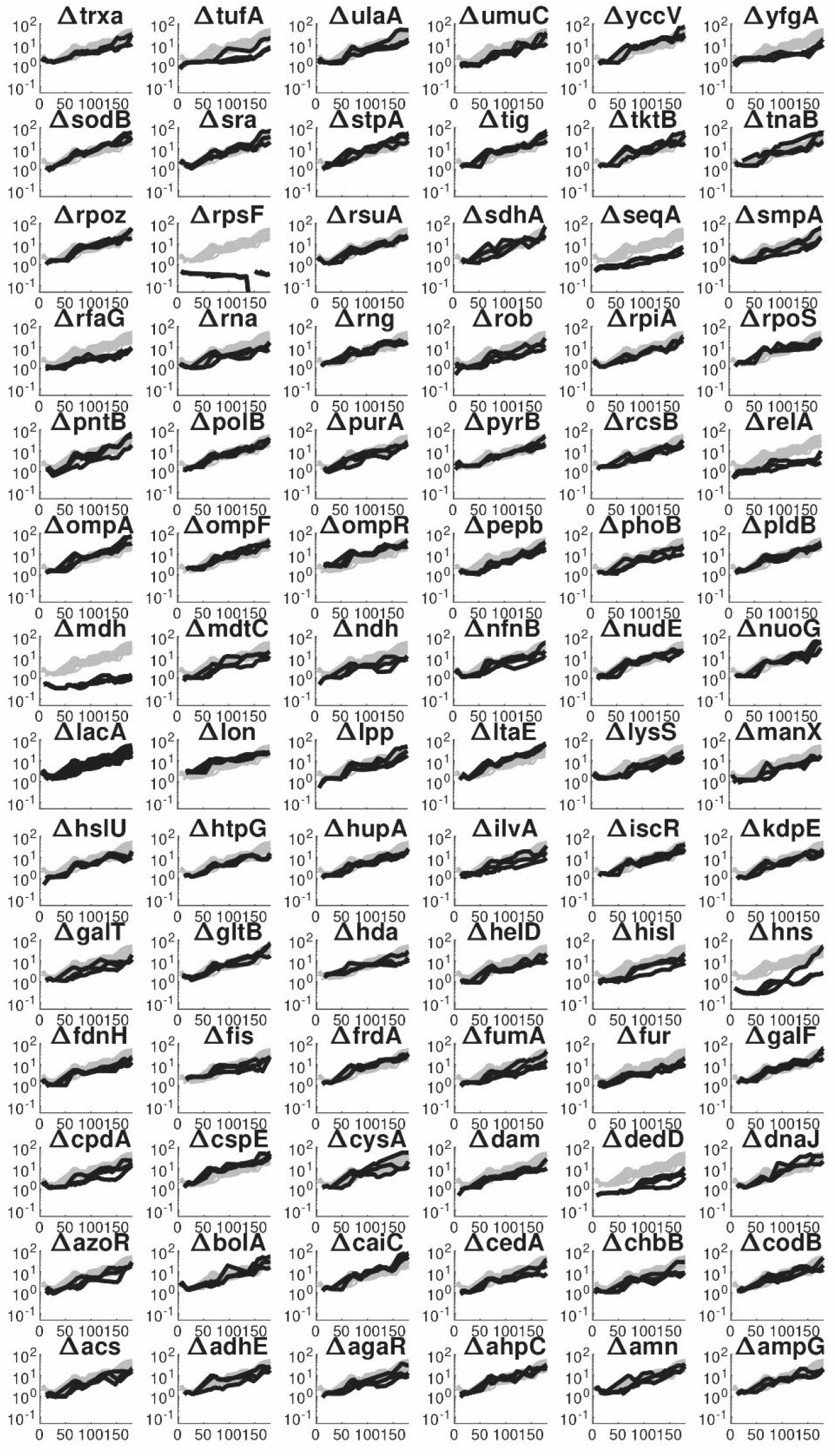
Resistance increases over time in chloramphenicol for additional ancestral strains. Resistance (as measured by the antibiotic concentration in the well) over time in chloramphenicol, but for all strains for which more than two replicates were done successfully (i.e. no cross-contamination was found).

**Table S1.**
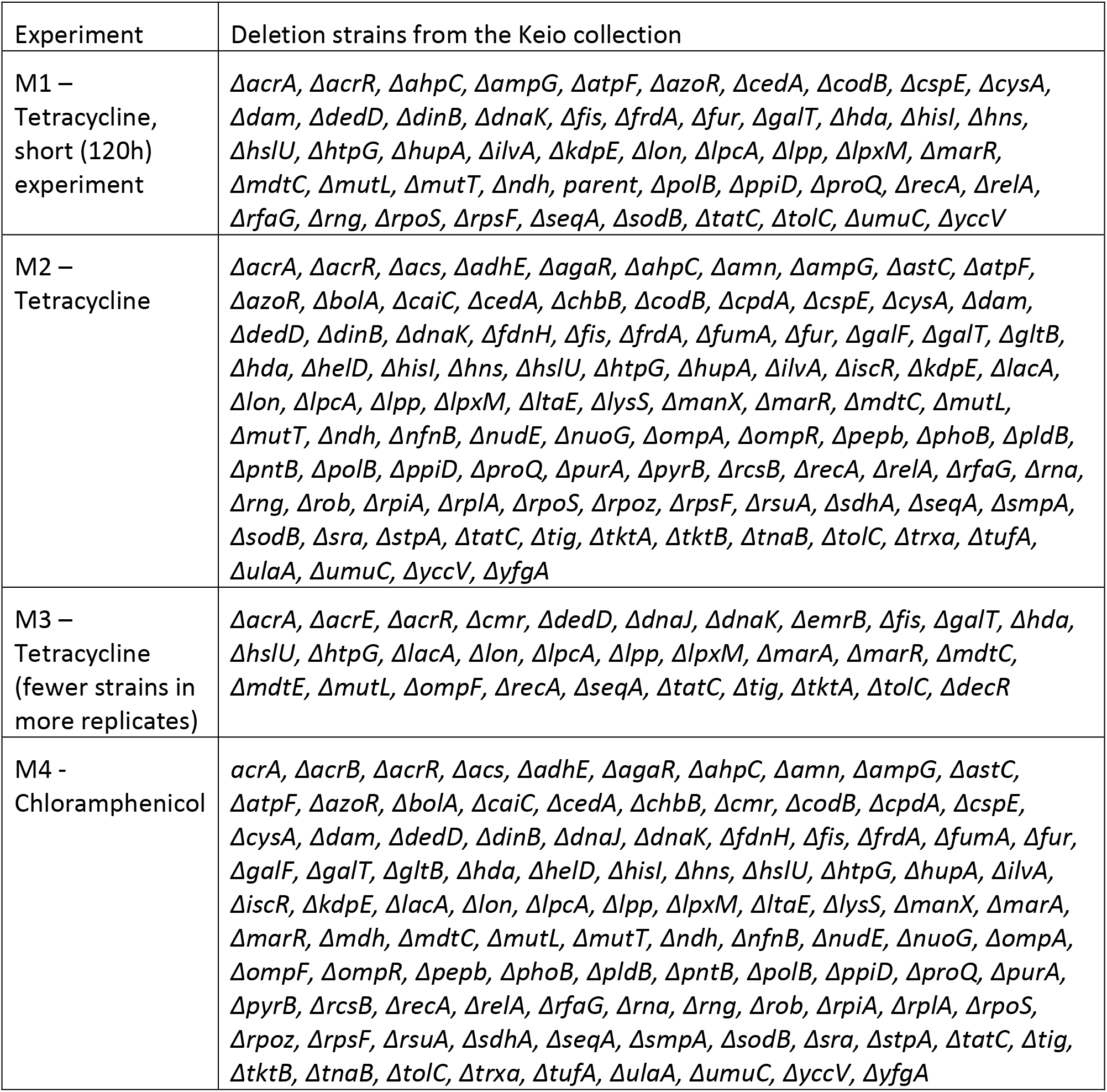
Lists of strains used in evolution experiments.

**Table S1.** Differences between instances of the automatized evolution experiments.

|  | M1 | M2 | M3 | M4 |
| --- | --- | --- | --- | --- |
| Antibiotic | Tetracycline | Tetracycline | Tetracycline | Chloramphenicol |
| Number of different strains | 51 | 99 | 31 | 104 |
| Replicates per strain | 3 | 3 | 8-12 | 3 |
| Control strain | Keio parent | $\Delta lacA$ | $\Delta lacA$ | $\Delta lacA$ |
| Replicates of control | 10 | 22 | 11 | 23 |
| Medium | LB | LB+50µg/ml kanamycin | LB+50µg/ml kanamycin | LB+50µg/ml kanamycin |

**Table S2.** Estimates of the resistance contributions of the 5 most common mutations based on linear regression model.

|  | Estimate | Standard error | t-statistic | p-value |
| --- | --- | --- | --- | --- |
| b0 | 1.348 | 0.198 | 6.82 | $2.55 \times 10^{-10}$ |
| b1 (*marR*) | 1.154 | 0.180 | 6.42 | $2.00 \times 10^{-9}$ |
| b2 (*lon*) | 1.144 | 0.180 | 6.35 | $2.81 \times 10^{-9}$ |
| b4 (pumps) | 0.2657 | 0.197 | 1.35 | 0.180 |
| b5 (*acrR*) | 0.3587 | 0.197 | 1.82 | 0.071 |
| b6 (*decR*) | 0.8096 | 0.181 | 4.48 | $1.55 \times 10^{-5}$ |

**Table S3.**
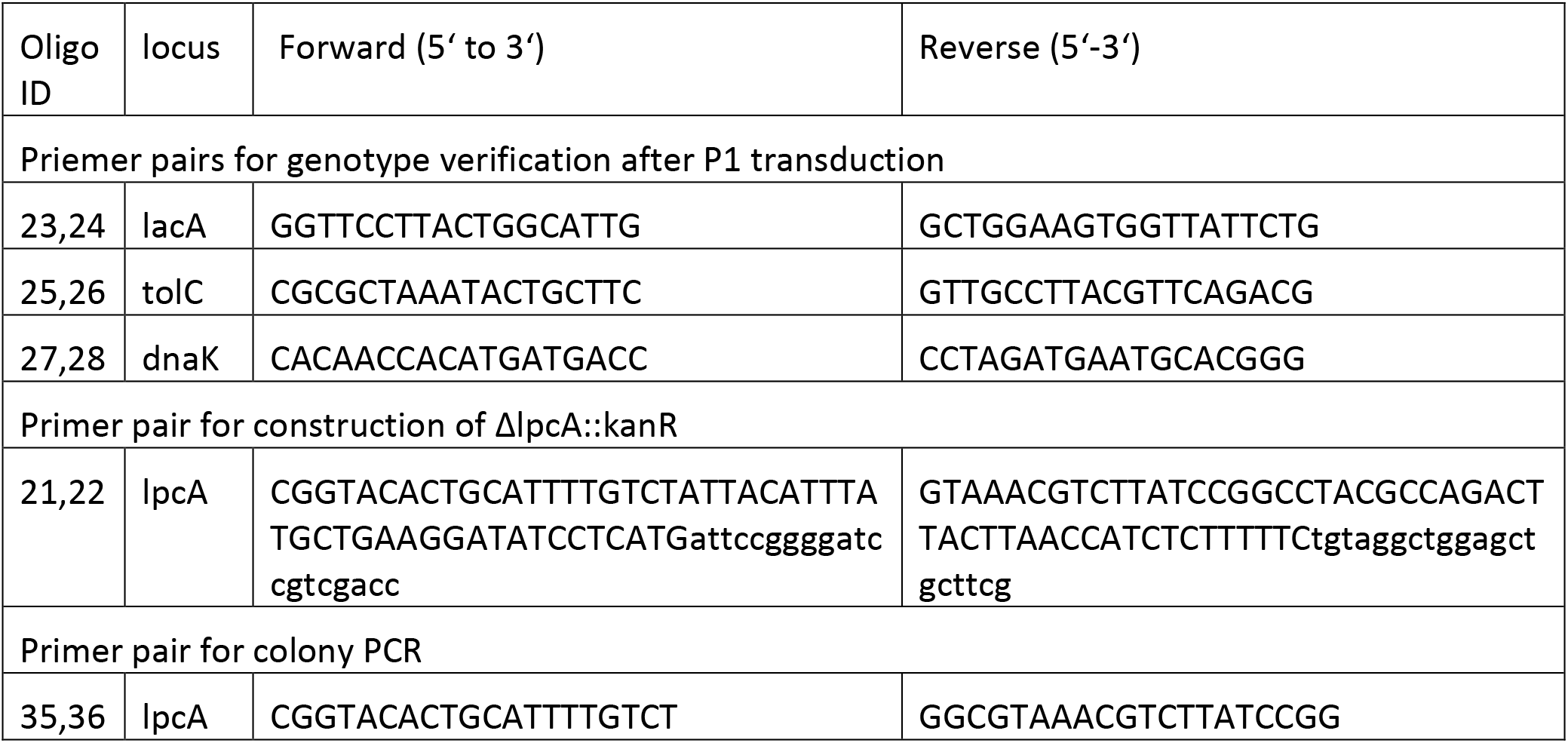
Primer pairs used for construction and verification of double deletion strains.

#### Data S1. (separate file)

Descriptions of all fixed mutations found in ancestral and evolved clones.

